# Early Blood Metabolome Remodeling Reveals Metabolic Signatures of Hypoxic-Ischemic Encephalopathy

**DOI:** 10.64898/2026.04.01.715878

**Authors:** Ashish Panigrahi, Neha Yadav, Swarupa Panda, Abinashi Sabysachi Sethi, Santosh Kumar Panda, Nirmal Kumar Mohakud, Vivek Tiwari

## Abstract

Hypoxic-ischemic encephalopathy (HIE) is a neonatal brain injury in which a definitive diagnosis within the first 6 hours of life is essential for initiating therapeutic hypothermia and improving neurological outcomes. However, early clinical evaluation and currently available biomarkers lack quantitative specificity during this narrow therapeutic window. We are providing insights into how rapid disturbances in systemic metabolites that regulate cerebral energy metabolism, neurotransmitter cycling, redox balance, and membrane integrity generate a definitive biochemical signature of HIE immediately after birth.

We performed quantitative blood-based 1H NMR metabolomics on collected blood samples within ∼1 hour of birth from 81 neonates (HIE, n = 42; non-HIE, n = 39), under optimized handling conditions to preserve metabolic integrity. Metabolite analysis revealed a distinct HIE-associated profile characterized by elevated lactate, alanine, succinate, glutamate, taurine, glycine, choline, and pyroglutamate, alongside significant depletion of glucose and glutamine compared with non-HIE controls (p < 0.05). These coordinated metabolic shifts reflect impaired mitochondrial respiration, enhanced anaerobic glycolysis, excitotoxic amino acid accumulation, altered membrane phospholipid turnover, and oxidative stress.

Multivariate analysis demonstrated clear separation between groups (PLS-DA accuracy = 0.83, R²Y = 0.46, Q² = 0.82), with glutamine, lactate, glutamate, and pyroglutamate as key discriminators. Pathway enrichment highlighted perturbations in glycolysis, the glucose-alanine cycle, glutamate-glutamine metabolism, Warburg effect like metabolic reprogramming, and redox homeostasis. Integration into supervised machine-learning models (Random Forest, XGBoost, SVM, KNN) achieved strong diagnostic performance (AUC = 0.97 ± 0.03; sensitivity ≈ 87%). Collectively, this minimally invasive NMR-to-machine-learning framework enables early, mechanistically grounded risk stratification of neonatal HIE within the therapeutic window.

## 1. Introduction

Impaired cerebral oxygen delivery and blood flow during the perinatal period underlie perinatal asphyxia and the subsequent development of hypoxic-ischemic encephalopathy (HIE), a neonatal brain injury associated with substantial risks of epilepsy, cerebral palsy, long-term neurodevelopmental deficits, and mortality ^1–4^. Therapeutic hypothermia is the only approved, definitive, and reliable neuroprotective intervention ^5^, which needs clinical confirmation of HIE within 6 hours after birth, hence is limited by the absence of definitive early quantitative signatures for timely intervention ^6–7^. Clinical scoring systems, neuroimaging, electroencephalography, and established circulating biomarkers predominantly reflect secondary or evolving consequences of hypoxic-ischemic stress and often lack sensitivity during the immediate postnatal period. Conventional approaches, such as APGAR scoring, are qualitative and prone to inter-individual readability, hence lack the precision in the immediate clinical confirmation of HIE, while established biomarkers, such as S100B, neuron-specific enolase (NSE), and creatine kinase-BB (CK-BB), exhibit delayed expression and suboptimal specificity ^8^. The transition from adequate oxygenation to hypoxia-ischemia may involve alterations in cerebral energy metabolism and be accompanied by coordinated changes in mitochondrial function, neurotransmitter cycling, redox balance, and membrane integrity. Quantitative assessment of these metabolic features, together with conventional clinical and imaging indicators, may improve characterization of HIE during the early postnatal period when neuroprotective strategies are most effective.

A panel-based, quantitative metabolomic strategy may overcome these limitations by capturing coordinated metabolic dyshomeostasis rather than singular biochemical deviations. Absolute quantification of metabolites central to brain energy metabolism, neurotransmitter cycling, redox homeostasis, and membrane integrity enables biologically interpretable comparisons across individuals and cohorts. When integrated with supervised machine-learning approaches, such data may reveal structured metabolic patterns associated with HIE, supporting a more precise and mechanistically informed framework for early diagnosis and risk stratification ^4, 9–12^. This study leverages integrated quantitative 1H-NMR plasma metabolomics with machine-learning analysis to generate a dynamic, multi-panel metabolic signature from blood collected within the first hours after birth, revealing early metabolic dyshomeostasis associated with neonatal HIE. Our findings present a concomitant decrease in brain energy neurochemicals such as glutamate-glutamine, glucose along with elevated levels of lactate, glutamate, alanine, succinate, taurine, choline and glycine in HIE compared to non-HIE controls. Machine-learning classifiers demonstrated high discriminatory performance (AUC up to 0.90), confirming that integrated metabolite quantification provides a precise, minimally invasive, and clinically translatable platform for early HIE diagnosis. This metabolic-ML framework offers a definitive strategy for rapid neonatal risk stratification and timely neuroprotective intervention.

## 2. Materials and Methods

### 2.1 Subject Selection and Demographic Details

Blood samples were collected from 105 neonates, of which 81 were included for 1H-NMR-based metabolomic analysis. Four samples were excluded due to clotting, and three due to insufficient plasma volume. An additional 13 samples were excluded from the final analysis owing to improper storage conditions and mishandling, while the remaining four were excluded due to excessive intervals between blood collection and plasma isolation. 42 Neonates with low APGAR scores (5.2 ± 1.2 at 5 minutes) were diagnosed with perinatal/birth-asphyxia labelled as probable cases of hypoxic-ischemic encephalopathy (HIE), based on clinical presentation, neurological signs, and medical records by clinicians. Blood samples of 39 neonates with high and consistent APGAR scores (9.7 ± 0.3 at 5 minutes), indicating healthy postnatal adaptation used as non-HIE/control. Non-HIE/control subjects were delivered without signs of maternal or fetal distress. They demonstrated normal respiration, intact neonatal reflexes, and stable physiological responses at birth. Collected samples from subjects were full-term infants (≥ 37 weeks gestation), ensuring exclusion of prematurity-related metabolic variability. Subject recruitment, eligibility screening, and application of inclusion/exclusion criteria were carried out under clinical supervision by clinicians. Demographic parameters such as gender, birth weight, and APGAR scores were recorded, and details are summarized in **Table 1**.

**Table 1:**
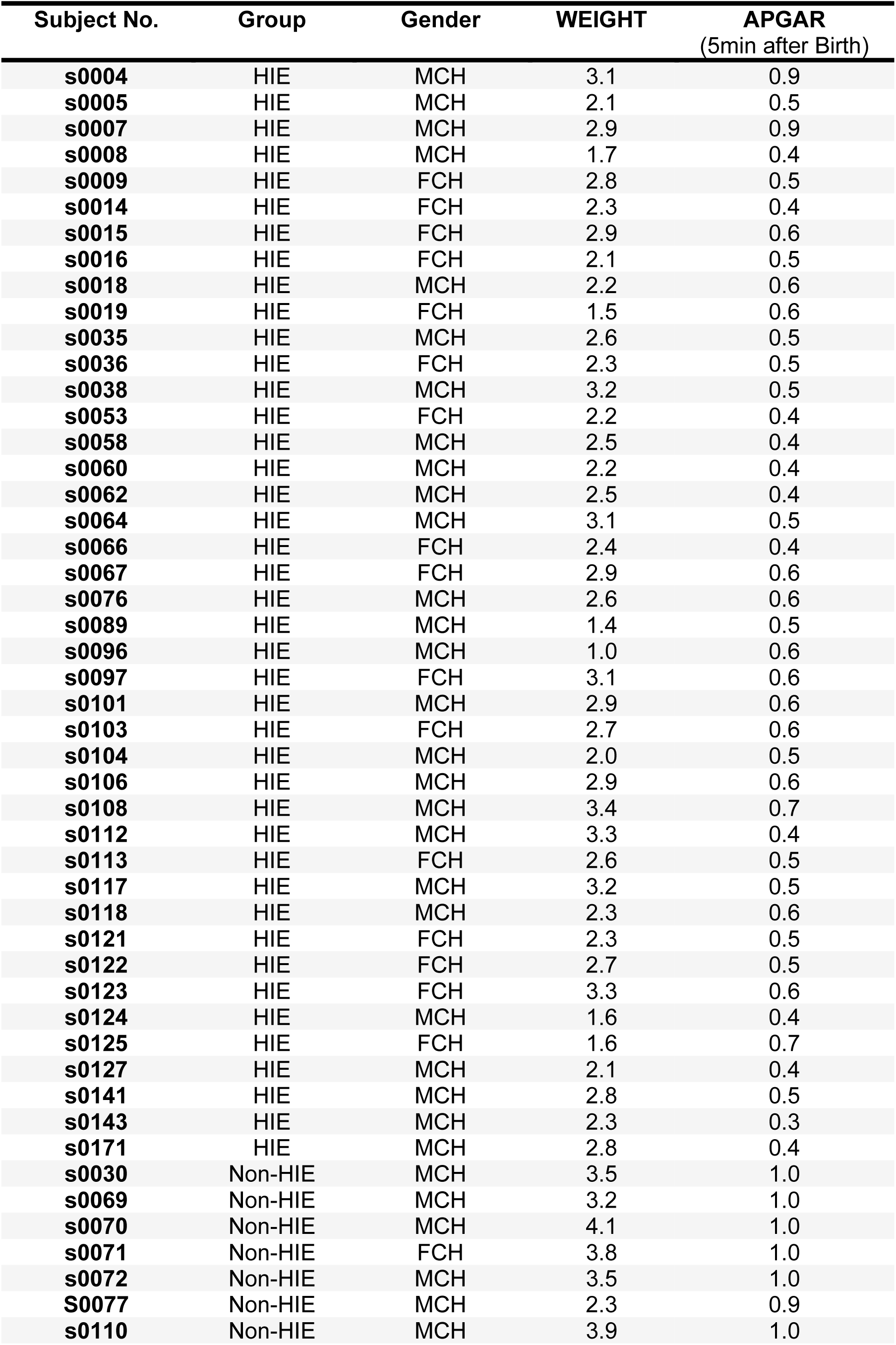
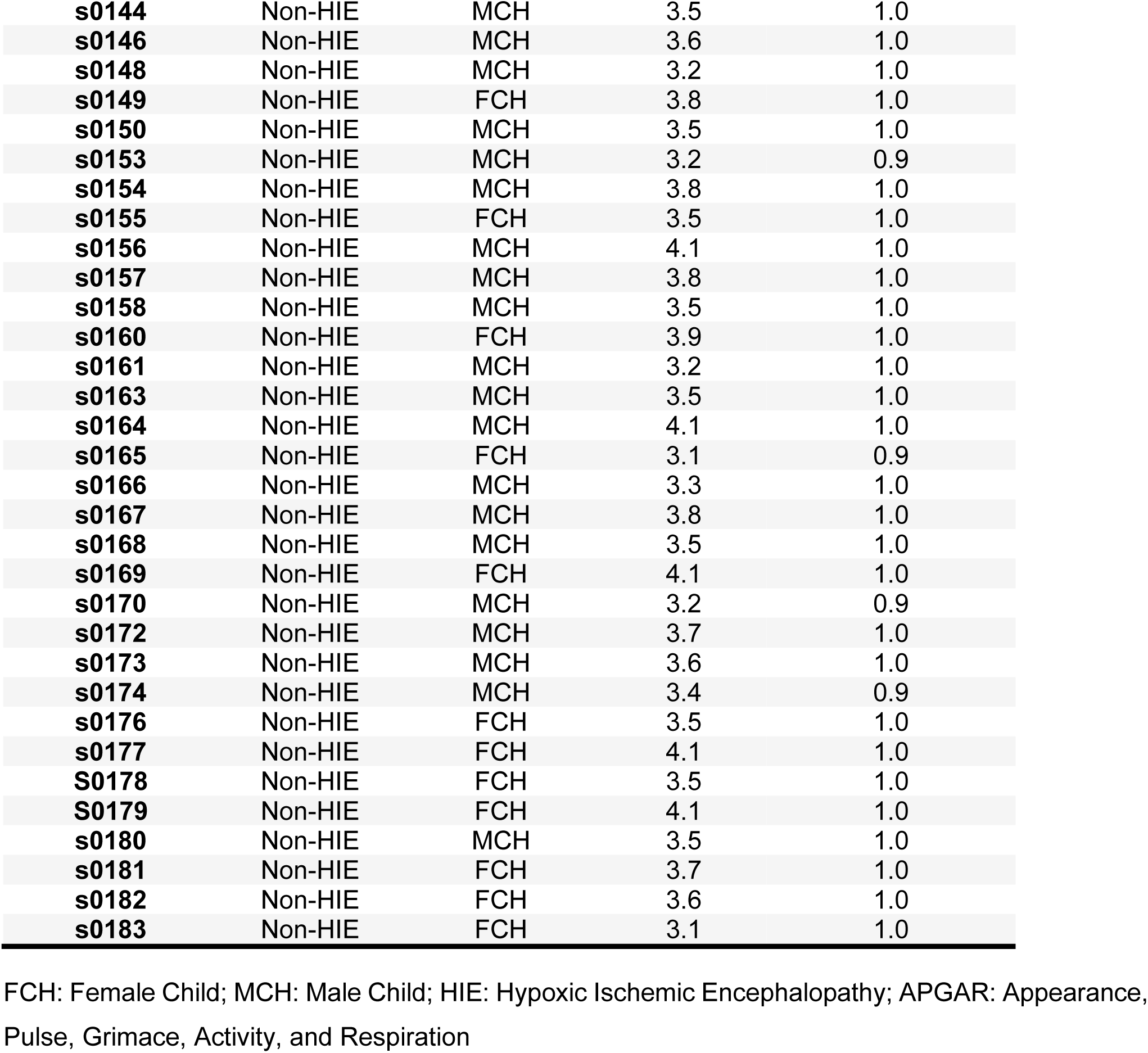
Demographic details of collected neonatal samples across 3 different medical colleges and hospitals of Odisha, India. Blood samples were collected from Birth/Perinatal Asphyxia neonates labelled as probable cases of HIE (n=42) and non-HIE (n=39).

| Subject No. | Group | Gender | WEIGHT | APGAR<br>(5min after Birth) |
| --- | --- | --- | --- | --- |
| s0004 | HIE | MCH | 3.1 | 0.9 |
| s0005 | HIE | MCH | 2.1 | 0.5 |
| s0007 | HIE | MCH | 2.9 | 0.9 |
| s0008 | HIE | MCH | 1.7 | 0.4 |
| s0009 | HIE | FCH | 2.8 | 0.5 |
| s0014 | HIE | FCH | 2.3 | 0.4 |
| s0015 | HIE | FCH | 2.9 | 0.6 |
| s0016 | HIE | FCH | 2.1 | 0.5 |
| s0018 | HIE | MCH | 2.2 | 0.6 |
| s0019 | HIE | FCH | 1.5 | 0.6 |
| s0035 | HIE | MCH | 2.6 | 0.5 |
| s0036 | HIE | FCH | 2.3 | 0.5 |
| s0038 | HIE | MCH | 3.2 | 0.5 |
| s0053 | HIE | FCH | 2.2 | 0.4 |
| s0058 | HIE | MCH | 2.5 | 0.4 |
| s0060 | HIE | MCH | 2.2 | 0.4 |
| s0062 | HIE | MCH | 2.5 | 0.4 |
| s0064 | HIE | MCH | 3.1 | 0.5 |
| s0066 | HIE | FCH | 2.4 | 0.4 |
| s0067 | HIE | FCH | 2.9 | 0.6 |
| s0076 | HIE | MCH | 2.6 | 0.6 |
| s0089 | HIE | MCH | 1.4 | 0.5 |
| s0096 | HIE | MCH | 1.0 | 0.6 |
| s0097 | HIE | FCH | 3.1 | 0.6 |
| s0101 | HIE | MCH | 2.9 | 0.6 |
| s0103 | HIE | FCH | 2.7 | 0.6 |
| s0104 | HIE | MCH | 2.0 | 0.5 |
| s0106 | HIE | MCH | 2.9 | 0.6 |
| s0108 | HIE | MCH | 3.4 | 0.7 |
| s0112 | HIE | MCH | 3.3 | 0.4 |
| s0113 | HIE | FCH | 2.6 | 0.5 |
| s0117 | HIE | MCH | 3.2 | 0.5 |
| s0118 | HIE | MCH | 2.3 | 0.6 |
| s0121 | HIE | FCH | 2.3 | 0.5 |
| s0122 | HIE | FCH | 2.7 | 0.5 |
| s0123 | HIE | FCH | 3.3 | 0.6 |
| s0124 | HIE | MCH | 1.6 | 0.4 |
| s0125 | HIE | FCH | 1.6 | 0.7 |
| s0127 | HIE | MCH | 2.1 | 0.4 |
| s0141 | HIE | MCH | 2.8 | 0.5 |
| s0143 | HIE | MCH | 2.3 | 0.3 |
| s0171 | HIE | MCH | 2.8 | 0.4 |
| s0030 | Non-HIE | MCH | 3.5 | 1.0 |
| s0069 | Non-HIE | MCH | 3.2 | 1.0 |
| s0070 | Non-HIE | MCH | 4.1 | 1.0 |
| s0071 | Non-HIE | FCH | 3.8 | 1.0 |
| s0072 | Non-HIE | MCH | 3.5 | 1.0 |
| S0077 | Non-HIE | MCH | 2.3 | 0.9 |
| s0110 | Non-HIE | MCH | 3.9 | 1.0 |
| **s0144** | Non-HIE | MCH | 3.5 | 1.0 |
| **s0146** | Non-HIE | MCH | 3.6 | 1.0 |
| **s0148** | Non-HIE | MCH | 3.2 | 1.0 |
| **s0149** | Non-HIE | FCH | 3.8 | 1.0 |
| **s0150** | Non-HIE | MCH | 3.5 | 1.0 |
| **s0153** | Non-HIE | MCH | 3.2 | 0.9 |
| **s0154** | Non-HIE | MCH | 3.8 | 1.0 |
| **s0155** | Non-HIE | FCH | 3.5 | 1.0 |
| **s0156** | Non-HIE | MCH | 4.1 | 1.0 |
| **s0157** | Non-HIE | MCH | 3.8 | 1.0 |
| **s0158** | Non-HIE | MCH | 3.5 | 1.0 |
| **s0160** | Non-HIE | FCH | 3.9 | 1.0 |
| **s0161** | Non-HIE | MCH | 3.2 | 1.0 |
| **s0163** | Non-HIE | MCH | 3.5 | 1.0 |
| **s0164** | Non-HIE | MCH | 4.1 | 1.0 |
| **s0165** | Non-HIE | FCH | 3.1 | 0.9 |
| **s0166** | Non-HIE | MCH | 3.3 | 1.0 |
| **s0167** | Non-HIE | MCH | 3.8 | 1.0 |
| **s0168** | Non-HIE | MCH | 3.5 | 1.0 |
| **s0169** | Non-HIE | FCH | 4.1 | 1.0 |
| **s0170** | Non-HIE | MCH | 3.2 | 0.9 |
| **s0172** | Non-HIE | MCH | 3.7 | 1.0 |
| **s0173** | Non-HIE | MCH | 3.6 | 1.0 |
| **s0174** | Non-HIE | MCH | 3.4 | 0.9 |
| **s0176** | Non-HIE | FCH | 3.5 | 1.0 |
| **s0177** | Non-HIE | FCH | 4.1 | 1.0 |
| **S0178** | Non-HIE | FCH | 3.5 | 1.0 |
| **S0179** | Non-HIE | FCH | 4.1 | 1.0 |
| **s0180** | Non-HIE | MCH | 3.5 | 1.0 |
| **s0181** | Non-HIE | FCH | 3.7 | 1.0 |
| **s0182** | Non-HIE | FCH | 3.6 | 1.0 |
| **s0183** | Non-HIE | FCH | 3.1 | 1.0 |

FCH: Female Child; MCH: Male Child; HIE: Hypoxic Ischemic Encephalopathy; APGAR: Appearance, Pulse, Grimace, Activity, and Respiration

Clinical data indicated significantly lower APGAR scores and higher rates of perinatal complications in the HIE group with respect to non-HIE. This stringent stratification enabled specific characterization of metabolic alterations attributable to hypoxic-ischemic injury while minimizing confounding effects of gestational age or other perinatal factors.

### 2.2. Blood Collection, Storage, and Processing

The study was approved by the Institutional Ethics Committees of all three participating medical colleges and hospitals (MKCG Berhampur, SCB Cuttack, and KIMS Bhubaneswar) and from Indian Institute of Science Education and Research (IISER) Berhampur. Peripheral blood samples (∼1.0 mL) were obtained from 105 neonates into heparinized vacutainers, ensuring immediate anticoagulation and rigorous preservation of sample integrity for downstream analyses. Immediately after collection, blood tubes were placed on ice and transported to the processing laboratory to maintain metabolic stability. Plasma was isolated by centrifugation at 6000 rpm for 10 minutes at 4 °C under standardized conditions. The supernatant plasma was carefully aliquoted into pre-labelled cryovials and stored at −80 °C to preserve biochemical integrity, minimizing degradation of proteins, metabolites, and other bioactive molecules. These plasma aliquots were subsequently used for 1H NMR-based metabolomic analyses **(Figure 1).**

**Figure 1:**
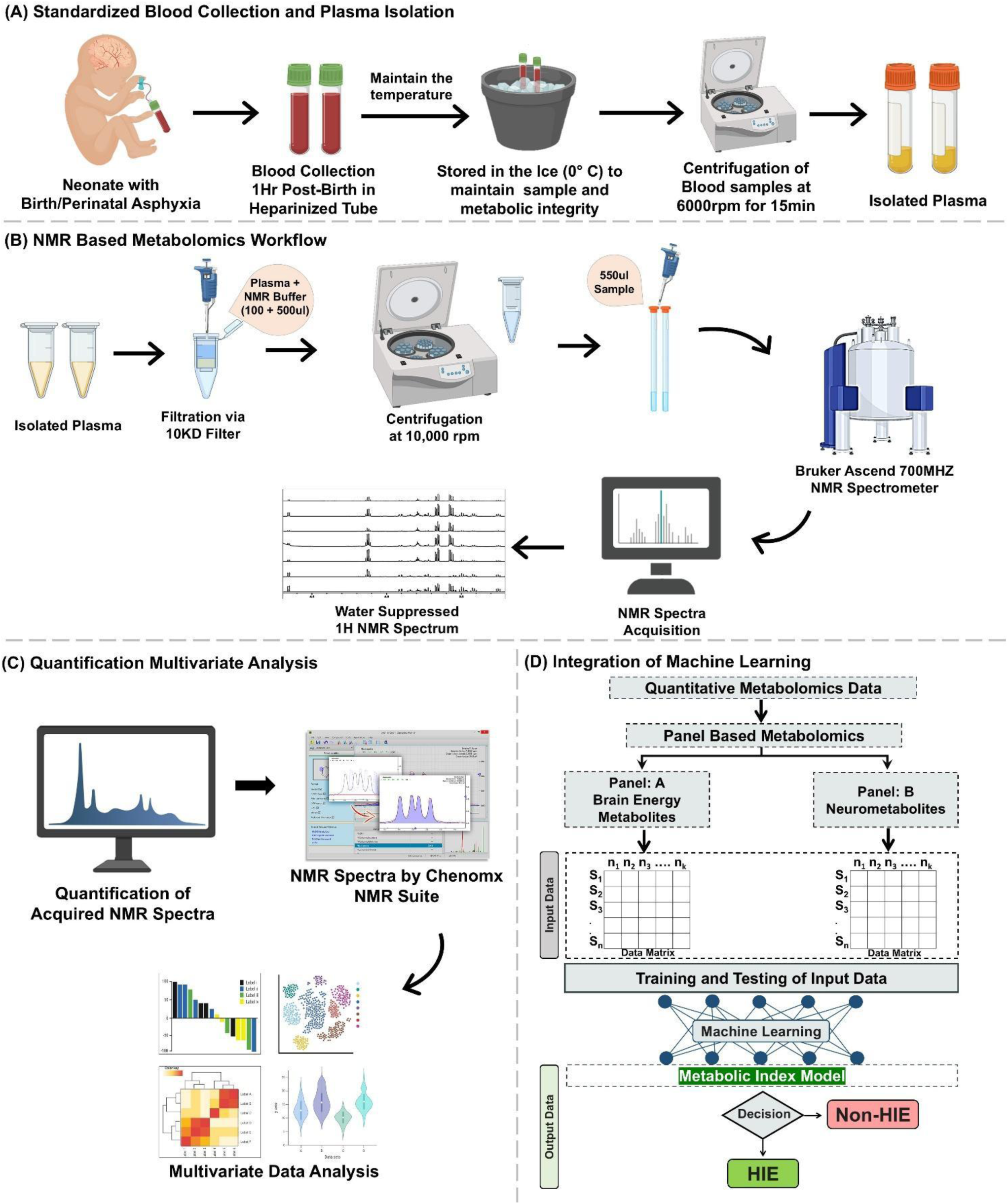
Integrated workflow for plasma metabolomics and machine learning–assisted biomarker discovery in neonatal hypoxic-ischemic encephalopathy (HIE). (A) Blood collection methodology: Neonatal blood samples are collected post-birth, temperature-controlled, and processed to isolate plasma. (B) Metabolomics pipeline: Isolated plasma undergoes 1H NMR-based metabolomic profiling following filtration and centrifugation steps, leading to water-suppressed 1H NMR spectral acquisition. (C) Data quantification and analysis: Acquired NMR spectra are quantified and subjected to comprehensive statistical and multivariate analysis. (D) Machine learning framework: Quantitative metabolomic and cytokine data are integrated in panel-based and global models using machine learning to generate predictive indices for HIE diagnosis. The figure is created using Biorender.com (https://BioRender.com/3j3xto9).

### 2.3. Blood-Based Metabolomics

Isolated plasma samples (100µl) were filtered using a pre-washed 10KDa centrifugal filter with NMR buffer. The NMR buffer consisted of PBS supplemented with D₂O and H₂O in a 20:80 ratio, along with 0.25 mM TSP (Trimethylsilyl propanoic acid) as a chemical shift indicator. The filtration step effectively removed high-molecular-weight proteins and macromolecules, thereby improving spectral quality. D₂O in the NMR buffer provided a stable deuterium lock in the NMR spectrophotometer, enhancing spectral resolution, while TSP served as an internal standard for chemical shift calibration and normalization. Processed plasma samples were subsequently used for 1H-NMR acquisition and analysed to identify and quantify metabolites. The resulting spectra enabled (a) metabolic profiling to assess the metabolite pool of birth-asphyxia-induced HIE and non-HIE/control neonates, and (b) the development of a precise quantitative blood-based metabolomics for HIE. Prior to use, the 10 KDa filters were pre-washed by adding 500 µL of distilled water followed by centrifugation at 6000 rpm for 10 min. This step was repeated four times to ensure complete removal of residual contaminants **(Figure 1)**.

### 2.4. NMR Spectroscopy and Data Acquisition Methods

Proton nuclear magnetic resonance (1H-NMR) spectroscopy was employed for metabolic profiling of plasma samples owing to its non-destructive nature, reproducibility, and capacity to simultaneously detect multiple low-molecular-weight metabolites. Spectra were acquired on Bruker Ascend 700MHz spectrophotometer using standardized protocols. To minimize water signal interference, pre-saturation with zero-gradient pulsed relaxation (ZGPR) sequences was applied. Deuterium oxide (D₂O) was included in the buffer to provide a field-frequency lock, and TSP served as a chemical shift reference. Automated frequency tuning and matching (ATMA) and gradient shimming were performed before acquisition to optimize magnetic field homogeneity. Spectral acquisition was carried out using 128 scans per sample with a receiver gain of 2.56Hz, ensuring adequate signal-to-noise ratio. Data were processed using TopSpin 4.0 and Chenomx V10, and chemical shifts were referenced to TSP at 0.0 ppm.

### 2.5. Data Processing and Metabolite Profiling

The acquired spectra were processed using TopSpin 4.0. Free induction decay (FID) signals were multiplied by an exponential function with a 0.5 Hz line-broadening factor and subjected to Fourier transformation (‘ft’ command) to improve the signal-to-noise ratio. The resulting spectra were manually calibrated to the TSP reference peak at 0.00 ppm, followed by zero-order and first-order phase correction. Metabolite quantification was performed using the Chenomx NMR Suite v.10. FID files were imported into the Chenomx processor module, baseline-corrected using the Whittaker spline function, and recalibrated to the chemical shift indicator. The processed spectra were then analysed in the Chenomx profiler module by matching metabolite cluster peaks to the software’s reference library. All the spectral concentrations were normalised by internal reference (TSP). Once peak alignment and normalizations were optimized, metabolite concentrations were calculated and exported in millimolar (mM) units. The following metabolites were quantified: lactate, glutamine, succinate, NADH, glucose, creatine phosphate, glutathione, N-acetylaspartate, myo-inositol, ATP, 2-hydroxybutyrate, choline, glycerol, histamine, phenylalanine, glycine, valine, alanine, glutamate, acetate, pyruvate, creatine, tyrosine, taurine, and pyroglutamate **(Table S1)**.

### 2.6. Multivariate Statistical Analysis and Pathway Enrichment

The partial least-squares discriminant analysis (PLS-DA) is a supervised classification model employed to obtain a global overview of the dataset to identify discriminating metabolites between HIE and non-HIE. The optimal PLS-DA model was identified using five components and evaluated via 5-fold cross-validations with R^2^ (model fit) and Q^2^ (predictive ability) indicating that the observed separation was not driven by random variation. Also, the metabolic loading for each PLS component was analysed to observe the contribution of each metabolite towards the separation. To evaluate metabolic differences between groups, volcano plots were generated to provide a combined visualization of fold-change magnitude and statistical significance for each metabolite. This approach allowed simultaneous identification of highly altered and statistically relevant metabolic features. In parallel, one-way analysis of variance (ANOVA)-based feature selection was performed to determine metabolites that significantly contributed to intergroup variability. The ANOVA model ranked features based on F-statistics and associated p-values, ensuring that only metabolites demonstrating robust group-dependent variation were retained for downstream analysis.

Metabolic pathway enrichment analysis was performed using the SMPDB-based pathway module in MetaboAnalyst 6.0. Significant metabolites identified in the differential analysis were mapped onto curated biochemical pathways to determine biologically relevant perturbations and metabolic system-level alterations. Pathways were ranked based on enrichment scores and pathway impact values to prioritize those most strongly affected in the disease state. To further characterize metabolic interactions and regulatory patterns, correlation analysis was conducted using Pearson’s correlation coefficients. The resulting correlation matrix was visualized as a hierarchically clustered heatmap to facilitate the identification of metabolite co-regulation networks. This approach enabled the detection of distinct metabolic modules spanning energy metabolism, amino acid biosynthesis and degradation, and oxidative redox balance. Together, these analyses provided an integrated systems-level view of metabolic dysregulation and highlighted functionally interconnected biochemical processes associated with the condition under investigation **(Figure 1).**

### 2.7. Machine learning models for discriminating HIE and non-HIE

To evaluate the diagnostic potential of metabolomic profiles, multiple supervised machine learning (ML) classifiers were used, including, XGBoost, Random Forest, Support Vector Machine (SVM), and KNN approach integrating model predictions. Data were randomly partitioned into training (80%) and testing (20%) sets, with all models optimized using 5-fold cross-validation to ensure robustness and prevent overfitting. Model performance was quantified using receiver operating characteristic (ROC) analysis and the area under the ROC curve (AUC), averaged across all cross-validation folds to assess classification stability and generalizability. These analyses quantified both the magnitude and direction of each metabolite’s contribution toward HIE prediction, providing biological insight into metabolic alterations underlying classification outcomes. Additionally, classifiers were independently trained using targeted metabolic panels including brain energy metabolites and neurometabolites to benchmark predictive performance against the integrated, full-feature ensemble model. Model precision, recall, specificity, and balanced accuracy were computed to evaluate discriminatory reliability, and confusion matrices were generated to visualize classification consistency across HIE and non-HIE **(Figure 1)**.

## 3. Results

### 3.1. Blood Based Metabolomics: Optimal Conditions for Metabolic Estimations

Optimization of storage condition, temperature and duration of the blood sample for precise metabolite estimation using 1H-NMR revealed that when blood samples were kept at room temperature (∼25°C), marked alterations in metabolite profiles were observed over time, wherein glucose levels decreased (−0.34t + 2.49) over the time, accompanied by a steady rise in lactate concentration (0.55t + 0.38), indicating ongoing glycolytic activity in the stored samples **(Figure 2A).** Representative 1H-NMR spectra of isolated plasma samples illustrating trends of pronounced reduction in glucose resonance with a concomitant elevation in lactic acid across the different storage time points (0 - 8 hours) **(Figure 2A).** Conversely, when blood samples were placed in ice immediately after collection, to maintain the metabolic integrity and minimise metabolic degradation. Under these conditions, glucose (−0.03t + 2.26) and lactate (0.07t + 0.81) concentrations remained largely constant even after different storage time points (0 - 8 hours) **(Figure 2B).** The corresponding 1H NMR spectra demonstrated minimal signal variation in glucose and lactate across time points, confirming enhanced metabolite preservation **(Figure 2B).** The concentration (mM) of metabolites were estimated for the two different storage and processing conditions **(Table S1).** Collectively, these findings emphasize that handling the plasma sample at subzero temperature following blood withdrawal is critical to preserve physiological metabolic integrity and minimize internal and external artefactual changes to reliably estimate the neurochemicals and metabolites.

**Figure 2:**
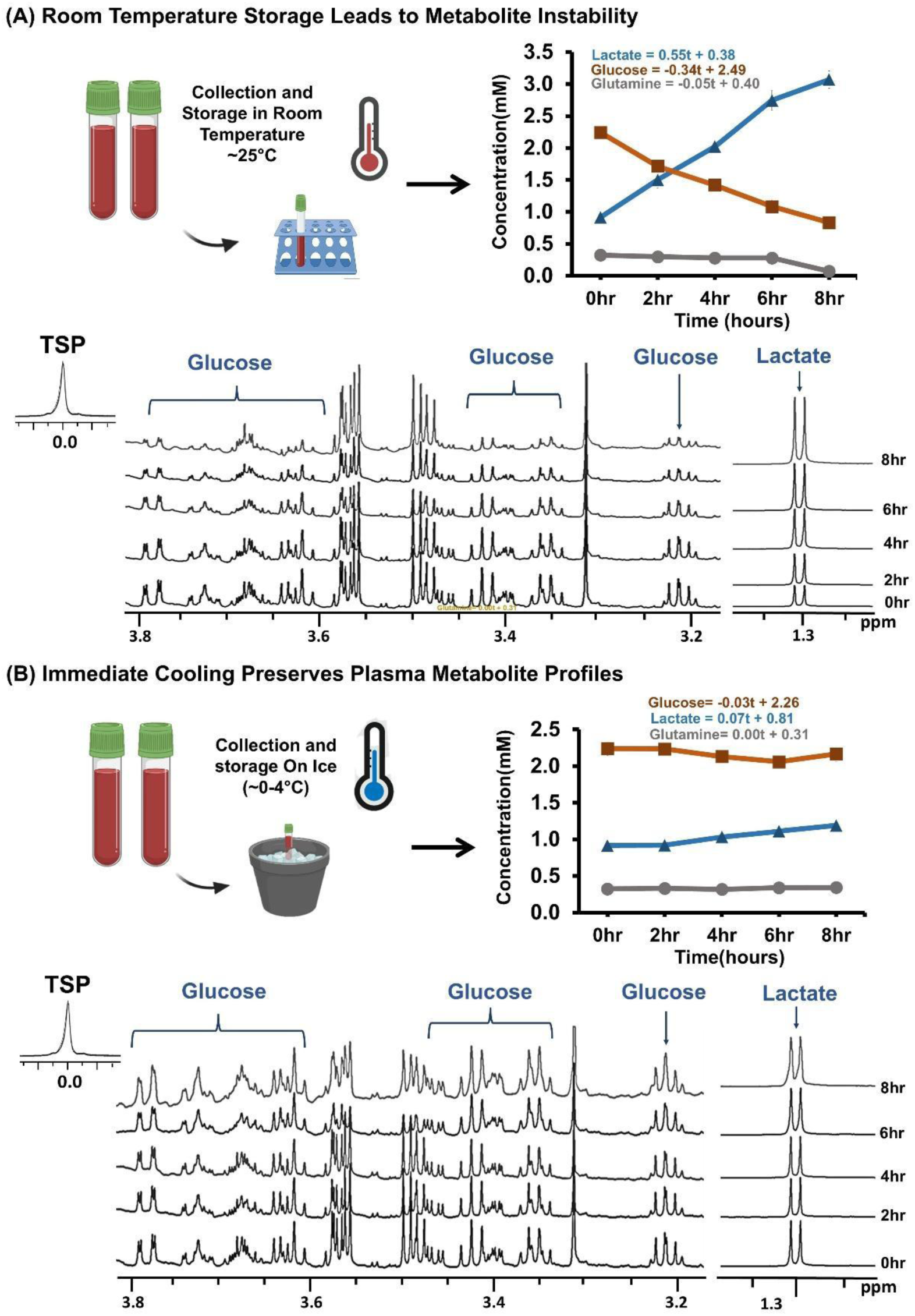
Influence of blood storage conditions on plasma metabolite stability for NMR-based metabolomics analysis. (A) Blood samples stored at room temperature (∼25°C) exhibit marked, time-dependent alterations in metabolite concentrations, with lactate increasing, glucose decreasing, and glutamine remaining relatively stable over an 8-hour period, as evidenced by both quantitative plots and shifts in NMR spectra. (B) Immediate cooling and storage on ice (∼0-4°C) preserves metabolic integrity, resulting in minimal fluctuations in lactate, glucose, and glutamine concentrations, with consistent 1H NMR spectral profiles throughout the 8-hour storage interval.

### 3.2. NMR-Based Plasma Metabolomics Reveals Dynamic Neurochemical and Energy Alterations

Highly resolved plasma metabolic spectra were obtained from neonates identified as plausible HIE (n = 42) and non-HIE (n = 39) neonates **(Figure 3A).** Estimation of metabolite concentration revealed marked and statistically significant concentration differences between HIE and non-HIE neonates (*F = 24.80, p = 2.26E-04*). Quantitative measurements of glycolytic and TCA cycle intermediates revealed elevated concentrations of lactic acid (HIE: 3.8 ± 1.5; non-HIE: 2.19 ± 0.84 mM, *p = 4.41E-07*), alanine (HIE: 0.25 ± 0.16; non-HIE: 0.17 ± 0.09 mM, *p = 2.04E-02*) and succinate (HIE: 0.015 ± 0.02; non-HIE: 0.006 ± 0.008 mM, *p = 4.25E-02*), but accompanied with reduction in glucose (HIE: 0.5 ± 1.0; non-HIE: 1.3 ± 1.1 mM, *p = 2.92E-03*) **(Figure 3B).**

**Figure 3:**
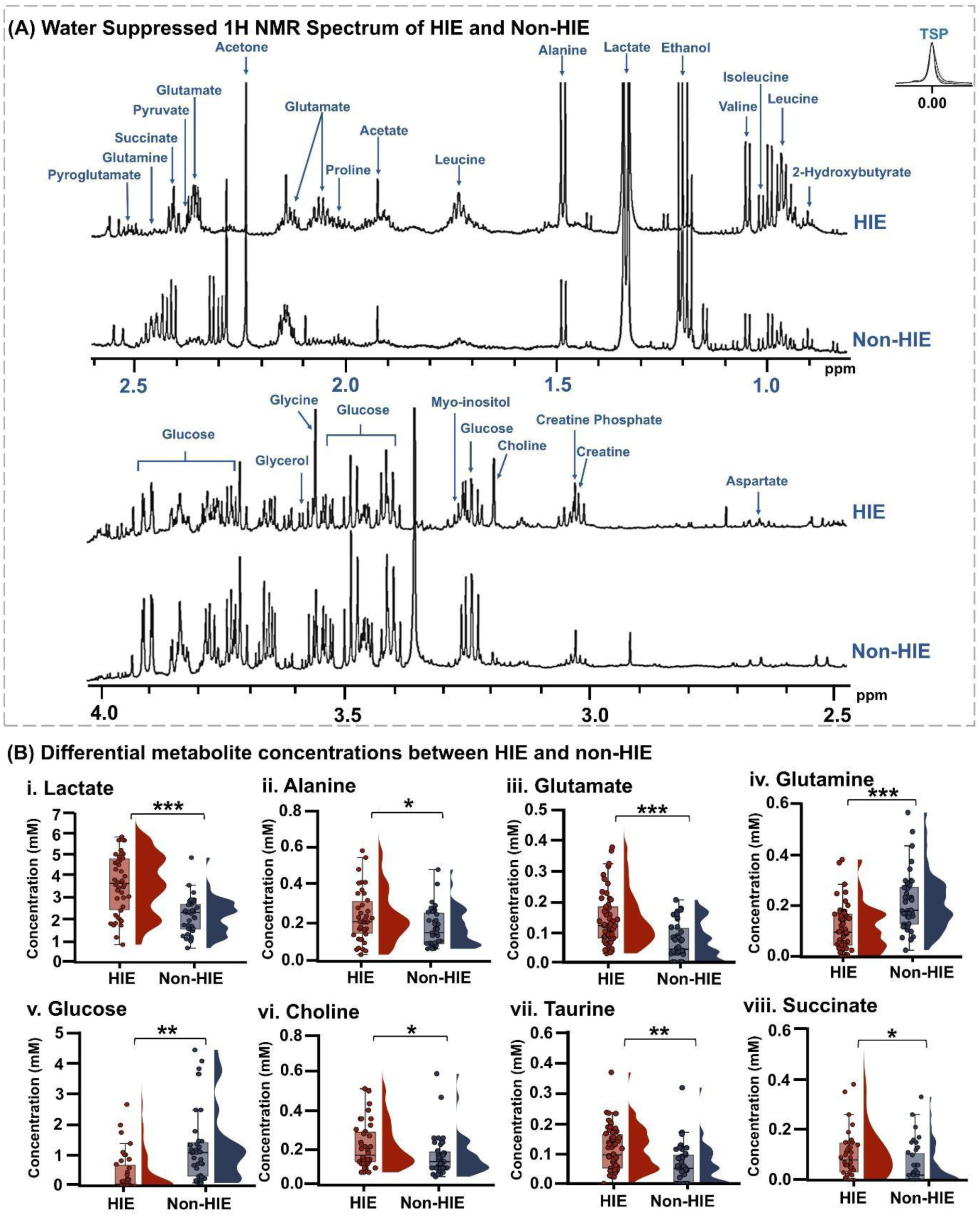
Comparative plasma-based metabolic profiling highlights distinct metabolic signatures using 1H-NMR. (A) Representative water-suppressed 1H NMR spectra display annotated resonances for major metabolites in HIE and non-HIE plasma samples, illustrating differences in metabolite abundance and spectral profiles. (B) Box plots summarize quantitative concentration differences for select metabolites, with significantly elevated or reduced levels of lactate, alanine, glycine, choline, glucose, succinate, glutamine, and glutamate in HIE compared to non-HIE neonates. Statistical significance is indicated as follows: * p < 0.05, ** p < 0.01, *** p < 0.001.

Wherein, HIE neonates displayed pronounced elevations in neurometabolites, reflecting altered neurometabolic homeostasis such as glutamate (HIE: 0.13 ± 0.1; non-HIE: 0.064 ± 0.06 mM, *p = 7.11E-04*), taurine (HIE: 0.12 ± 0.08; non-HIE: 0.07 ± 0.07 mM, *p = 1.01E-02*) was accompanied with reduction in glutamine (HIE: 0.09 ± 0.07; non-HIE: 0.3 ± 0.1 mM, *p = 3.62E-08*) **(Figure S1)**. Cellularity index depictive marker choline shows significant elevation (HIE: 0.02 ± 0.01; non-HIE: 0.015 ± 0.009 mM, *p = 3.31E-02*). The concentration values of the complete set of metabolites (n=25) were quantified, as shown in **Figure S1**. Moreover, these quantitative findings delineate a distinct metabolic signature characteristic of HIE marked by disrupted energy metabolites, excitotoxic amino acid accumulation, mitochondrial dysfunction, and impaired redox state reflecting the biochemical complexity underpinning hypoxia-ischemia induced neuropathology.

### 3.3. Multivariate Analyses of Metabolic Data

Supervised PLS-DA achieved clear class separation between significantly altered metabolites in HIE and non-HIE neonates **(Figure 4B).** Model reliability was evaluated using R² and Q² values derived from five-fold cross-validation. The optimized five-component PLS-DA model demonstrated strong classification performance (accuracy = 0.83), with good model fit with class labels (R²Y = 0.46), and high cross-validated predictive ability (Q² = 0.82), indicating a satisfactory discrimination between HIE and non-HIE despite biological heterogeneity **(Figure 4B; Table S2).** Discriminatory metabolites contribute to group separation in PLS1 and PLS2 according to the metabolic loading **(Figure 4B and D).** These identified metabolites are primarily mapped to perturbed biochemical pathways associated with impaired energy metabolism, mitochondrial dysfunction, and excitotoxic amino acid dysregulation.

**Figure 4:**
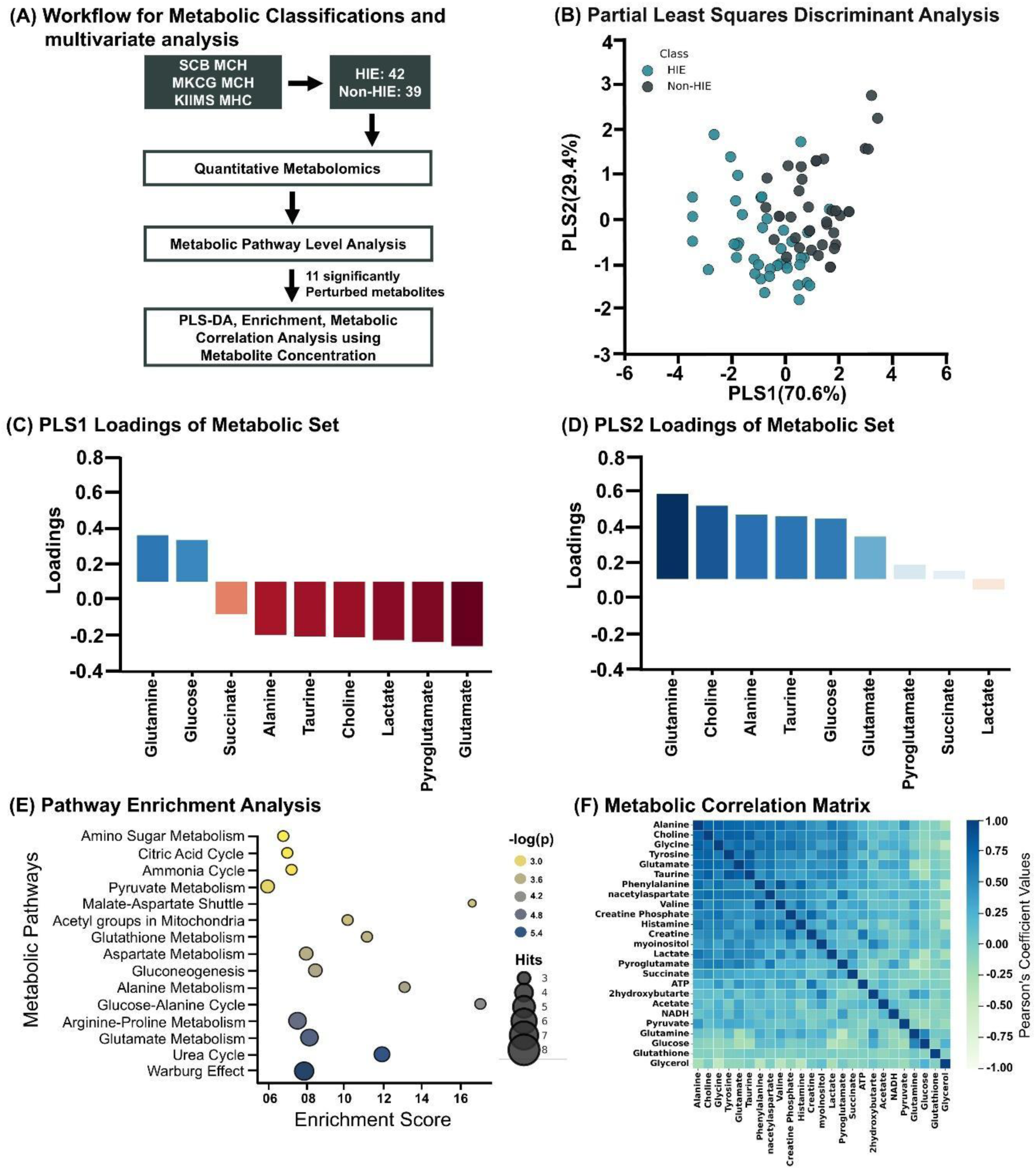
This figure summarizes integrative metabolomics and statistical analyses in neonatal samples distinguishing HIE from non-HIE subjects. (A) The workflow depicts steps from cohort selection to quantitative metabolomics and multivariate analyses. (B) Partial Least Squares Discriminant Analysis (PLS-DA) score plot visualizes separation between HIE and non-HIE based on metabolomic profiles. (C) PLS1 loading plot identifies metabolites most influential in principal component 1 for discriminating sample classes. (D) PLS2 loading plot shows metabolites mainly driving variance in principal component 2. (E) Pathway enrichment analysis highlights key metabolic pathways perturbed in HIE, showing enrichment scores and statistical significance. (F) The metabolic correlation matrix reveals coordinated changes and correlations among quantified metabolites, clustering metabolic networks altered in HIE.

Volcano plot analysis revealed significant metabolic perturbations between HIE and non-HIE groups. Prominent differentially upregulated metabolites included glutamate (log₂FC = 1.026, *p = 8.06E-04*), succinate (log₂FC = 1.28, *p = 4.68E-02*), and pyroglutamate (log₂FC = 3.88, *p = 9.77E-05*) and downregulated metabolite glucose (log₂FC = −1.318, *p = 2.72E-03*) and glutamine (log₂FC = −1.279, *p = 1.50E-08*) **(Table S3).** These alterations reflect an increased glycolytic flux, excitotoxic amino acid accumulation, and disruption of the glutamine-glutamate cycle and metabolic stress responses in HIE neonates **(Figure S2A).**

Global analysis such as ANOVA reveals glutamine, lactate, glutamate and pyroglutamate are significant metabolites. Glutamine as the most significant discriminatory metabolite (p = 1.50E-08; F = 39.8 and t = 6.23), followed by lactate (p = 4.35E-07; F = 30.3 and t = −5.62), pyroglutamate (p = 9.77E-05; F = 16.9 and t = −4.25) and glutamate (p = 8.06E-04; F = 12.1 and t = −3.55) **(Table S4).** These findings point toward major dysregulation in glutamate-glutamine metabolism, oxidative stress pathways, and anaerobic glycolysis. To further interpret these metabolic disturbances, correlation and pathway enrichment analyses were conducted **(Figure S2B).**

### 3.4. Pathway Enrichment Analysis and Metabolic Correlation

Pathway enrichment analysis conducted using the MetaboAnalyst 6.0 platform revealed substantial perturbations across multiple biologically relevant metabolic pathways **(Figure 4E).** The most significantly enriched pathways included the Warburg effect, urea cycle, glutamate metabolism, arginine-proline metabolism, and the glucose-alanine cycle, each demonstrating strong statistical significance with -log₁₀(p) values ranging from ∼4.8 to 5.4. In parallel, notable alterations were also observed in gluconeogenesis, aspartate metabolism, and glutathione metabolism, further highlighting widespread dysregulation of energy homeostasis, nitrogen turnover, and cellular redox balance **(Figure 4E).** Together, these enriched pathways illustrate a coordinated metabolic reprogramming in HIE neonates, aligning with hallmarks of hypoxia-induced mitochondrial dysfunction, excitotoxicity, and oxidative stress. This systems-level metabolic disruption highlights the mechanistic relevance of the identified metabolites and supports their potential utility as candidate biomarkers of HIE progression and severity. A complete list of enrichment scores and corresponding -log₁₀(p) values for all significantly affected pathways is provided in **Table S5**.

Pearson’s correlation analysis highlights distinct metabolic co-regulation patterns, revealing well-defined biochemical modules associated with hypoxic injury. A prominent positively correlated cluster emerged among lactate, succinate, pyruvate, and ATP (r ≈ 0.47-0.66), indicating a tightly linked metabolic shift toward anaerobic glycolysis and disrupted mitochondrial oxidative phosphorylation **(Figure 4F).** Similarly, glucose, glutamine, and glycerol demonstrated coordinated behaviour, with weaker associations to glycolytic intermediates (r ≈ 0.25-0.43), suggesting partial metabolic compensation and altered substrate utilization under hypoxic stress **(Figure 4F)**. Additionally, a highly coordinated amino acid cluster comprising glutamate, alanine, glycine, taurine, tyrosine, and phenylalanine (r ≈ 0.65-0.85) was observed, suggesting synchronized regulation of neurotransmitter precursor pathways and increased amino acid turnover in response to hypoxic excitotoxicity **(Figure 4F)** Collectively, these correlation patterns delineate a structured and biologically meaningful metabolic network, highlighting coordinated impairment of energy metabolism, amino acid turnover, and neuronal integrity in HIE neonates.

### 3.5. Machine Learning Classifier Performance and Key Discriminatory Metabolites

Machine learning models integrating plasma metabolomic profiles demonstrated strong discrimination in distinguishing HIE from non-HIE neonates **(Figure 5).** The schematic workflow for the classification model is presented in **Figure 5A**. Balanced accuracy is around ∼88% of all the models, with XGB and Random Forest achieving the most consistent performance **(Figure 5B; Table S6)**. All tested models such as Random Forest, XGBoost, SVM, and KNN achieved high accuracy and balanced classification performance. Receiver operating characteristic (ROC) analyses revealed excellent model discrimination, with XGB and Random Forest exhibiting the highest mean (AUC = 0.97 ± 0.03) and (AUC = 0.97 ± 0.03) respectively, followed by KNN (AUC = 0.95 ± 0.04), and SVM (AUC = 0.90 ± 0.06) **(Figure 5C(III); Table S6)** and their confusion matrix **(Figure S3C).** Models showing minimal variation across all the folds emphasize the stability and efficacy of the model with ∼87% sensitivity. Two focused panels were designed from the 09 significantly perturbed metabolites identified in global analyses: Panel A; encompassing brain energy metabolites (glucose, lactate, succinate, and alanine), and Panel B, comprising neurometabolites (glutamine, glutamate, choline, and taurine). Each panel was separately evaluated using all 4 models **(Figure 5C I and II; Table S6)**. In the Brain energy metabolite panel, all classifiers demonstrated excellent predictive capacity, achieving high sensitivity up-to ∼83% across validation folds. Random Forest achieved the best overall performance (AUC = 0.90 ± 0.07), followed by KNN (AUC = 0.88 ± 0.07), XGBoost (AUC = 0.86 ± 0.07), and SVM (AUC = 0.77 ± 0.11) (**Figure 5C(I); Table S6)** and their confusion matrix **(Figure S3A).** Balanced accuracy analysis confirmed stable and reproducible classification efficiency, with mean accuracies consistently exceeding ∼79% **(Figure 5B).** In contrast, models trained on the neurometabolite panel exhibited slightly lower AUC-ROC values compared to brain energy metabolites but still substantial classification performance has been observed. KNN yielded the highest (AUC = 0.94 ± 0.07), followed by Random Forest (AUC = 0.92 ± 0.09), XGBoost (AUC = 0.89 ± 0.10) and SVM (AUC = 0.85 ± 0.11) **(Figure 5C(II); Table S6)** and their confusion matrix **(Figure S3B).** Balanced accuracy plots indicated moderate yet consistent accuracy ∼82%, and sensitivity in the neurometabolites panel is ∼82% demonstrating the biological relevance of neurometabolites for differentiation of HIE neonates **(Figure 5B).** Collectively, these findings highlight that both energy metabolism and neurotransmission related metabolite panels possess significant diagnostic value for HIE classification. These metabolic panels are distinctive in predicting HIE at the early stage of life and can be a prominent biomarker to diagnose HIE development in neonates.

**Figure 5:**
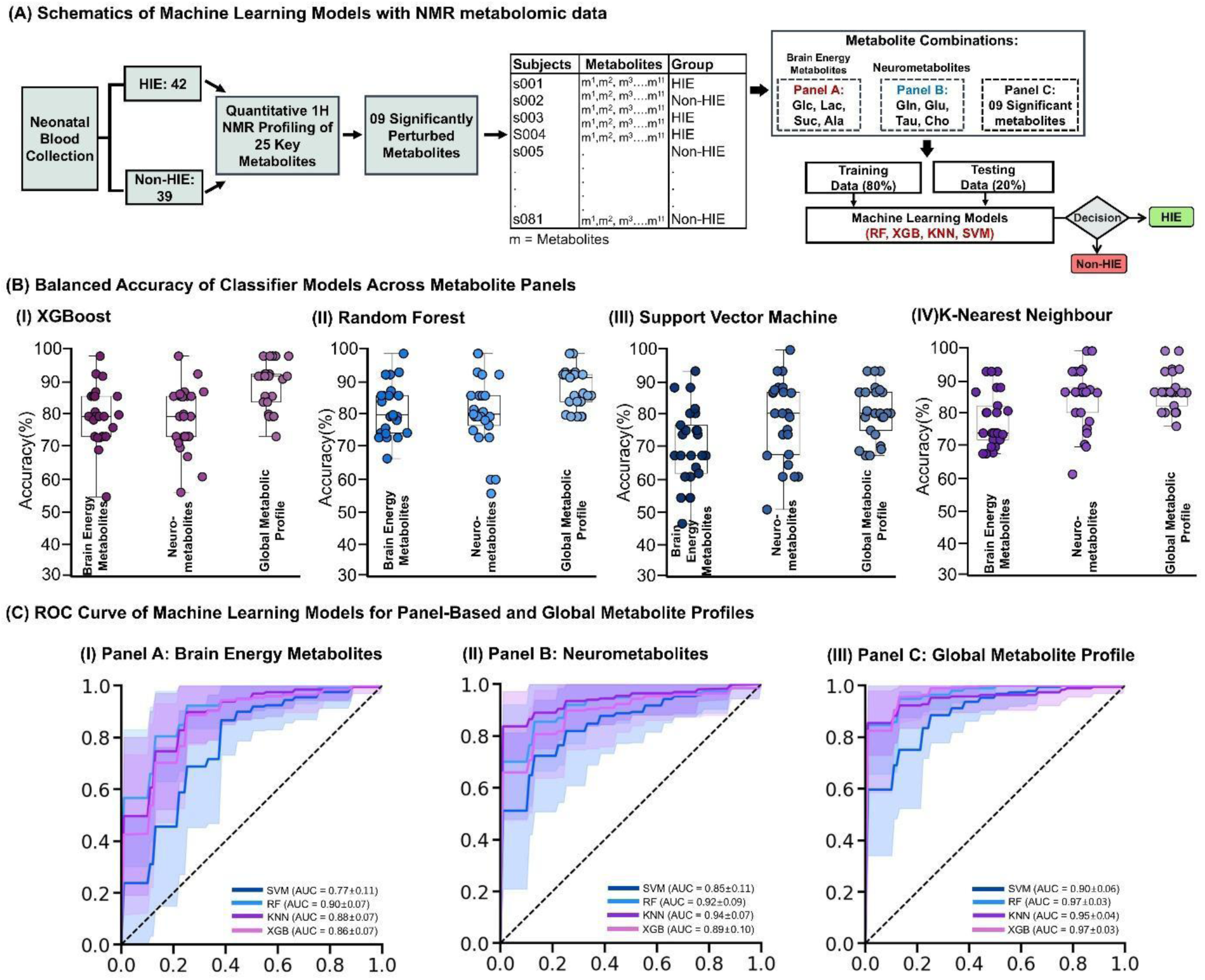
Overview and Performance of Machine Learning Models Using NMR based Quantitative Metabolomics Data in HIE Classification (A) Overview of study workflow utilizing neonatal blood metabolomic profiles for machine learning analysis. Following blood collection, 1H NMR profiling quantified 25 metabolites, from which 09 significantly altered metabolites were selected and organized into three panels based on HIE vs. non-HIE classification. Panel A: brain energy metabolites, Panel B: neurometabolites and Panel C: global metabolic profile. (B) Balanced accuracy of classifier performance is shown for XGBoost, Random Forest, Support Vector Machine (SVM), and k-Nearest Neighbour (KNN) across the three metabolite panels. (C) AUC-ROC curves for classifier models applied to each metabolite panel: Panel A (brain energy metabolites), Panel B (neurometabolites), and Panel C (global metabolite profile). Curves for SVM, RF, KNN, and XGB models are presented.

## 4. Discussion

Early diagnosis of HIE remains a major clinical challenge, as current diagnostic strategies do not adequately reflect the rapid and systemic metabolic reprogramming that follows hypoxic-ischemic injury. Despite evidence of extensive metabolic disturbances in urine, cord blood, and in-vivo models, these biochemical signatures lack definitive specificity for HIE and may be influenced by other neonatal conditions ^1, 13–14^, However, peripheral blood studies are more reliable and provides a practical window on global metabolic status of the neonate within hours after birth ^4, 15^. This study provides quantitative blood-based metabolomics coupled with multivariate statistics and machine-learning providing the first detailed description of the early postnatal plasma metabolome in HIE neonates within 1-hr of birth, revealing distinct metabolic fingerprints and potential biomarker candidates that differentiate HIE from non-HIE controls.

Pathway enrichment analysis demonstrated that the most strongly perturbed pathways cluster around brain energy metabolism, including glycolysis, the TCA cycle, pyruvate metabolism, the malate-aspartate shuttle, gluconeogenesis, and the Warburg-effect like reprogramming of energy flux **(Figure 4E).** Consistent with this, four core energy metabolites; lactate, succinate, glucose, and alanine showed marked and coordinated alterations at birth, indicating a switch from oxidative phosphorylation to anaerobic glycolysis with partial TCA cycle blockade **(Figure 6).** Elevated lactate and succinate reflect impaired mitochondrial respiration and accumulation of downstream TCA intermediates, whereas declining glucose suggests increased glycolytic demand to maintain ATP under hypoxia supported by in-vivo MRS ^16–17^. In parallel, alanine accumulation, linked to the glucose-alanine cycle, supports carbon and nitrogen shuttling between tissues and liver during anaerobic stress **(Figure 6).** These metabolic alterations in Panel A (glucose, lactate, alanine, succinate) define a coherent biochemical signature of impaired brain energy metabolism and clearly demonstrate the discriminative capacity of the machine learning model between HIE and non-HIE neonates (AUC: 0.85 ± 0.08) **(Figure 5C(I)).**

**Figure 6:**
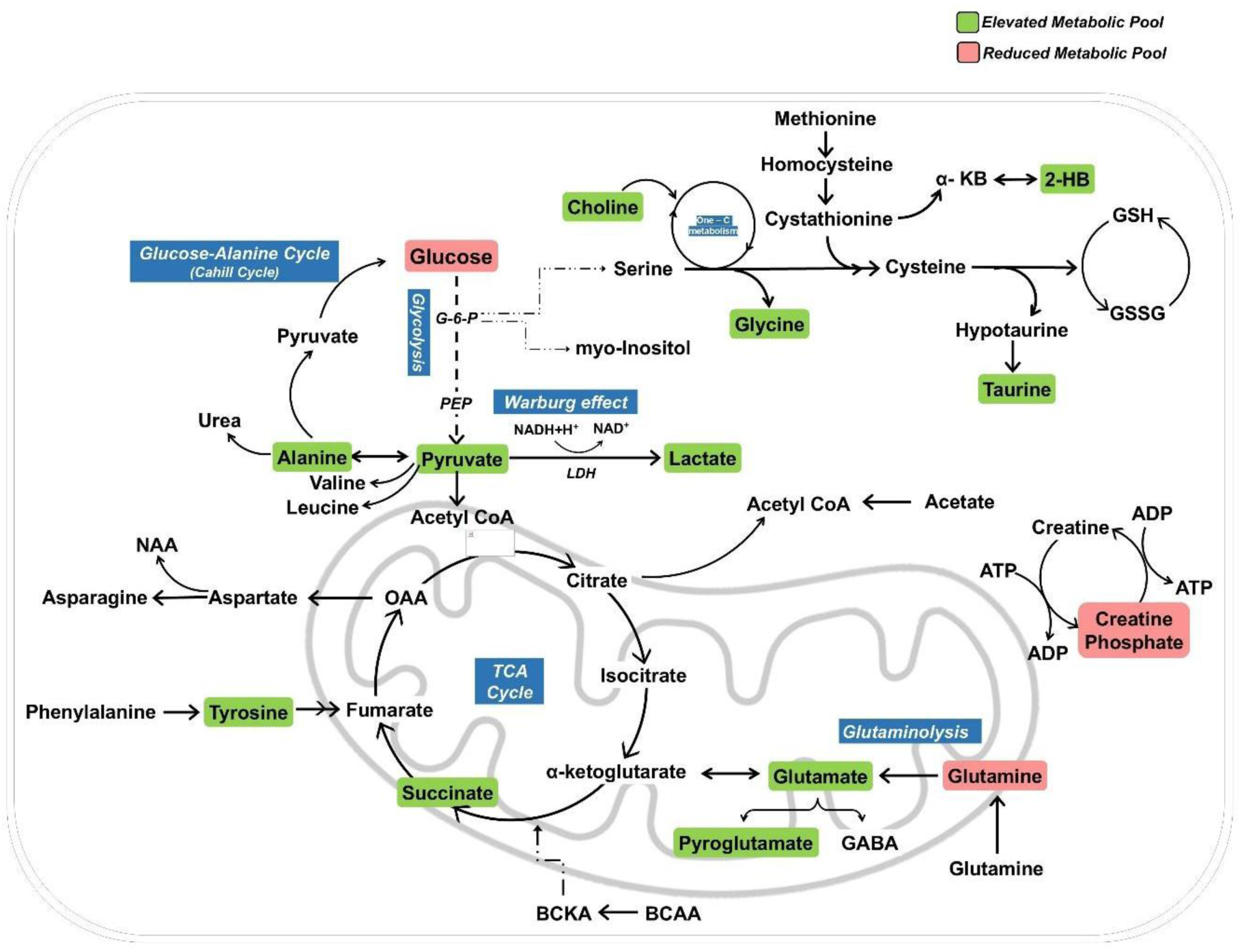
Integrated metabolic pathway map of HIE vs non-HIE neonates. The schematic highlights glycolysis, TCA cycle, Warburg effect, Glutaminolysis, one-carbon metabolism with elevated metabolites pool (green) or reduced metabolic pool (red). Elevated pools include lactate, alanine, succinate, glutamate, glycine, choline, taurine, pyruvate, 2-hydroxybutyrate, tyrosine, and pyroglutamate, while creatine phosphate and glutamine are reduced.

In addition to altered energy pathways, neurometabolites involved in neurotransmission and glial support such as glutamine, glutamate, choline, and taurine exhibited a highly conserved co-regulation pattern across correlation and enrichment analyses. This tightly coupled metabolic module reflects impaired neurometabolic homeostasis characteristic of HIE. Increased glutamate, together with altered glutamine-glutamate balance and enrichment of glutamate and aspartate metabolism pathways, points to excitotoxic signalling and impaired neurotransmitter recycling ^8, 17–18^. Elevations in glycine and taurine, both inhibitory or neuromodulatory amino acids, likely reflect compensatory responses to excitotoxic and oxidative stress, consistent with their roles in antioxidant defence, glutathione synthesis, and NMDA receptor modulation, and their strong positive correlations with other amino acids in the correlation matrix ^10, 19–20^. Wherein, choline cellularity index depictive marker shows significant alteration between HIE and non-HIE. That supports disruption of phosphatidylcholine (Kennedy) pathway activity and enhanced membrane phospholipid turnover, as reported in animal models of neonatal hypoxia where choline is similarly increased in plasma and urine ^21–22^. Also, elevated choline reflects convergent injury to neuronal membranes, synaptic integrity, and lipid metabolism, reinforcing its potential as an early marker of hypoxic-ischemic damage ^10, 20–22^. Collectively, these neurometabolic perturbations (Panel B: glutamine, glutamate, choline, taurine) form a distinct biochemical fingerprint indicative of impaired neuronal integrity and a disrupted excitatory-inhibitory balance in the neonatal brain. This coordinated profile not only reflects the pathophysiological consequences of hypoxic-ischemic injury but also reinforces the discriminatory strength of the machine-learning model in separating HIE from non-HIE neonates (AUC: 0.90 ± 0.09) **(Figure 5C-II).**

Redox homeostasis was profoundly affected in HIE, evidenced by a robust elevation of pyroglutamate relative to non-HIE controls **(Figure 3A; Figure S1).** As a central metabolite of the γ-glutamyl cycle, pyroglutamate serves as a biochemical proxy for glutathione turnover; thus, its accumulation indicates increased antioxidant demand and sustained oxidative pressure. This pattern aligns with previous evidence showing that hypoxia and critical neonatal illness accelerate glutathione depletion and trigger compensatory upregulation of γ-glutamyl intermediates ^23–24^. Notably, pyroglutamate, which clusters with energy-related and amino acid metabolites in the correlation matrix, demonstrates that redox disruption is not isolated but mechanistically intertwined with mitochondrial dysfunction, glutamate-glutamine cycling, and broader neurometabolic reprogramming. Collectively, these findings reveal the susceptibility of neonatal redox defences under hypoxic-ischemic burden and position pyroglutamate as a compelling candidate biomarker of glutathione pathway stress in HIE ^23–24^.

Integrating 09 significant metabolic signatures in Panel C (lactate, succinate, glucose, alanine, glutamine, glutamate, choline, taurine, and pyroglutamate) markedly improved classification performance, achieving a robust discriminative accuracy within the machine-learning framework (AUC: 0.94 ± 0.04) **(Figure 5C-III).** This panel-based multi-analyte strategy surpasses the diagnostic limitations of single metabolite markers and aligns with growing evidence that composite metabolic signatures offer superior clinical precision in HIE. By defining coordinated metabolic modules spanning brain energy pathways, neurotransmitter cycling, membrane turnover, and redox regulation, this panel provides mechanistic clarity on how hypoxia-ischemia disrupts interconnected biochemical processes in the neonatal brain at early hours of life. Collectively, these findings establish a biologically meaningful foundation for the development of metabolomics-informed diagnostic tools and future neuroprotective interventions in HIE.

## 5. Conclusion

This study establishes a robust, blood-based metabolomics and machine learning framework for early identification of potential biomarkers for neonatal HIE. Quantitative metabolomics data from peripheral plasma reveal a potential metabolic signature, characterized by elevated lactate, succinate, alanine, glutamate, taurine, choline, and pyroglutamate together with depleted glucose and glutamine, reflecting impaired mitochondrial respiration, disordered glycolytic-TCA coupling, and excitotoxic stress. Supervised multivariate and pathway analyses converged on profound perturbations in brain-energy metabolism, glutamate-glutamine cycling, and redox homeostasis, while panel-based machine learning classifiers leveraging these significantly altered metabolites achieved high diagnostic accuracy (AUC up to ∼0.90) for discriminating HIE from non-HIE neonates. Hence, these findings demonstrate that minimally invasive plasma metabolomic profiling, integrated with quantitative NMR and interpretable machine learning models, can yield clinically actionable early biomarkers for HIE and elucidate the underlying metabolic cascade that precedes overt neurological injury. This integrative approach offers a scalable platform for rapid neonatal risk stratification, optimization of the therapeutic hypothermia window, and future development of metabolically informed neuroprotective strategies.

## Abbreviations

HIE: Hypoxic Ischemic Encephalopathy
NMR: Nuclear magnetic resonance
FCH: Female Child
MCH: Male Child
APGAR: Appearance, Pulse, Grimace, Activity, and Respiration
AI-ML: Artificial Intelligence - Machine Learning
NSE: Neuron-Specific Enolase
CK-B0B: creatine kinase-BB
PBS: Phosphate Buffer
ZGPR: zero-gradient pulsed relaxation
D₂O: Deuterium oxide
TSP: Trimethylsilyl propionic acid
ATMA: Automated frequency tuning and matching
SVM: Support Vector Machine
KNN: K-nearest neighbour
ROC: Receiver Operating Characteristic
AUC: Area Under Curve

## Acknowledgments

The authors sincerely acknowledge the clinical collaborators and nursing staff of the three Medical Colleges and Hospitals in Odisha, India, for their invaluable support in sample collection and clinical assessment. We extend our gratitude to the technical staff at the IISER Berhampur NMR Facility for their assistance with NMR acquisition. Special thanks to the Odisha Science and Technology Department for funding and supporting this study. We express our deep appreciation to the parents and families who consented to participate, making this research possible. The authors would also like to thank Keith Hulsey for proofreading the article.

## Associated Content

## Data and Software Availability

NMR spectral preprocessing and metabolite profiling were performed using TopSpin 4.0 ([TopSpin | NMR Data Analysis | Bruker](https://www.bruker.com/en/products-and-solutions/mr/nmr-software/topspin.html)) and Chenomx NMR Suite v.10 ([Chenomx Inc | Metabolite Discovery and Measurement](https://www.chenomx.com/)). The graphical figures were created using BioRender ([Scientific Image and Illustration Software | BioRender](https://www.biorender.com/)).

## Author Contributions

**V.T.** - Conceptualization, original draft writing, literature review, figures and illustrations, NMR and metabolism discussions, machine-learning discussion, clinical challenges discussion, interpretation of clinical information, review and editing. **A.P.** - Original draft writing, figures and illustrations, NMR and metabolism discussions, machine-learning discussion, clinical challenges discussion, review and editing. **N.Y.** - Figures and illustrations, machine-learning discussion, review and editing. **S.P., A.S., S.K.P.** - Neonatal Blood sample collection, clinical assessment of participants, interpretation of clinical information, review and editing. **N.M.** - Conceptualization, neonatal blood sample collection, clinical assessment of participants, interpretation of clinical information, review and editing.

## Notes

The authors declare that there are no conflicts of interest related to this work. No financial, personal, or professional relationships that could influence the content of this review have been disclosed.

## Funding Source

This study is funded by the Department of Science & Technology, Government of Odisha, Grant Number: S&T/OD/BIO/BPR/010124/050.

## Data and Software Availability

All the codes are available in the private GitHub repository (https://github.com/nibr-lab/HIE-Manuscript), which will be made publicly available upon acceptance of the manuscript. In case of any clarifications, data can be directly provided upon request.

## Notes

### Competing Interest Statement

The authors have declared no competing interest.

