## Supplementary figures and tables for "Early Blood Metabolome Remodeling Reveals Metabolic Signatures of Hypoxic-Ischemic Encephalopathy"

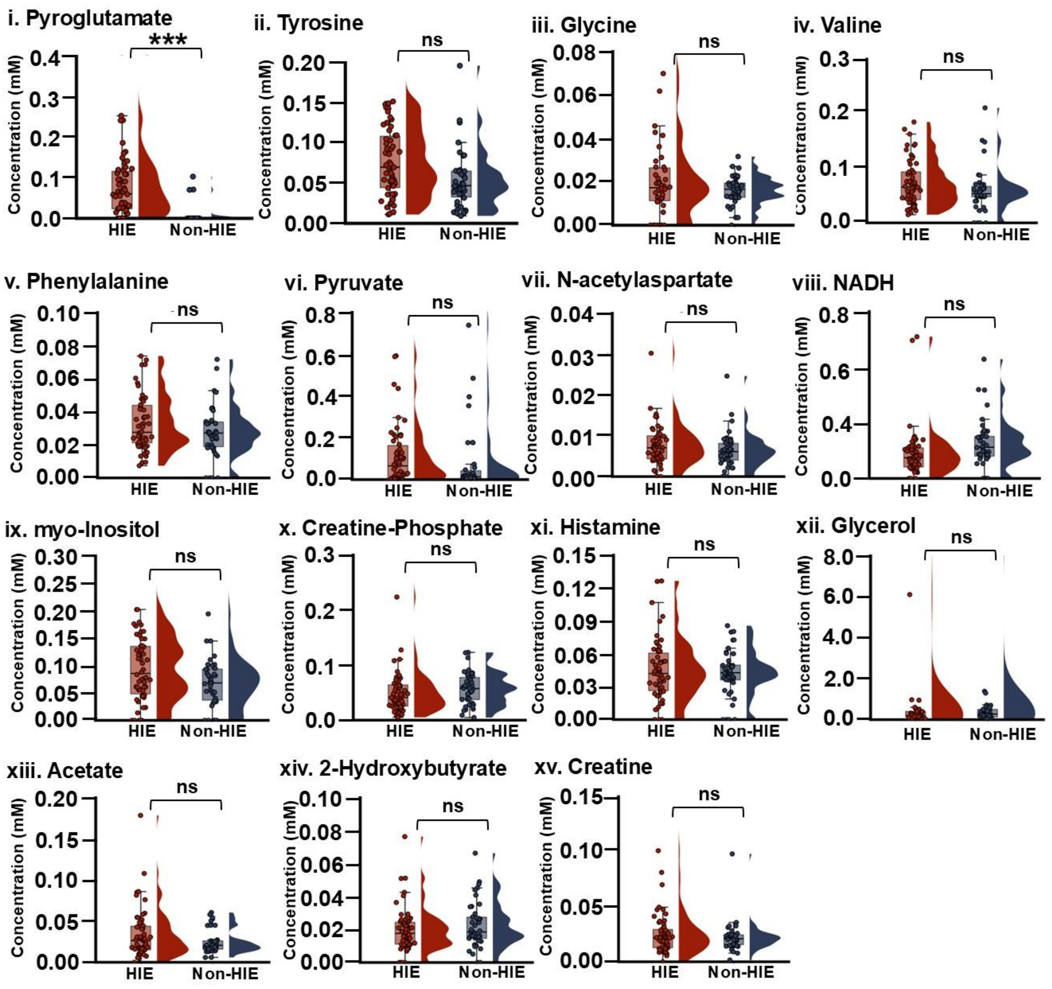


**Figure S1:** Raincloud plots illustrate the distribution of plasma concentrations for quantified key metabolites in neonates diagnosed with HIE and non-HIE. Statistical significance is indicated as follows: * $p < 0.05$, ** $p < 0.01$, *** $p < 0.001$, ns = not significant ($p \geq 0.05$).

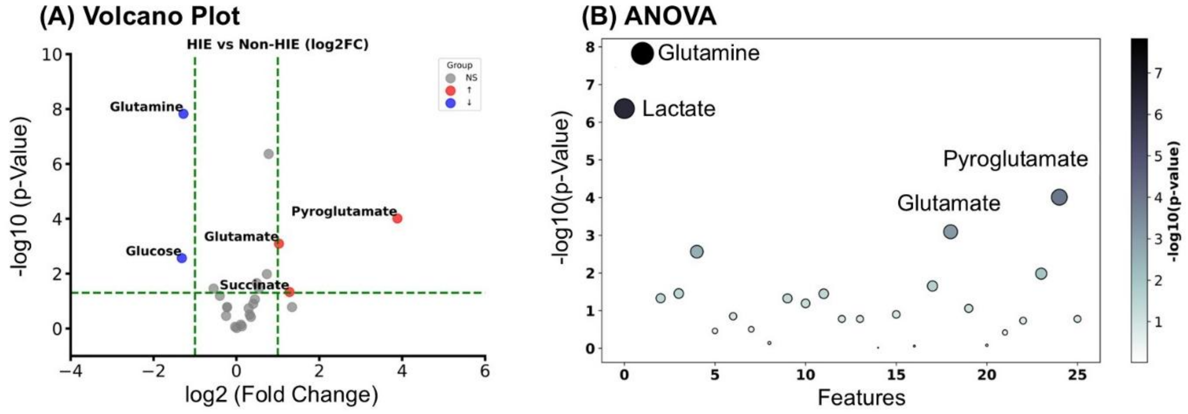


**Figure S2:** Identification of Significantly Altered Plasma Metabolites Distinguishing HIE from Non-HIE Neonates. (A) Volcano plot displaying the distribution of plasma metabolites based on log2 fold change and statistical significance (−log10(p-value)) between HIE and non-HIE neonatal groups. Each point represents a metabolite, colored by group (blue: upregulated in HIE, red: downregulated, grey: not significant). Lactate, glutamate, pyroglutamate, succinate, and glucose showed the most significant differences, with lactate and pyroglutamate markedly elevated in HIE. (B) Bubble plot of ANOVA (analysis of variance) results for plasma metabolites. Size and color intensity correspond to the strength of statistical significance (−log10(p-value)) for each metabolite. Lactate, glutamine, glutamate, and pyroglutamate exhibit the highest statistical significance, highlighting major metabolic perturbations in energy metabolism and amino acid turnover associated with HIE.

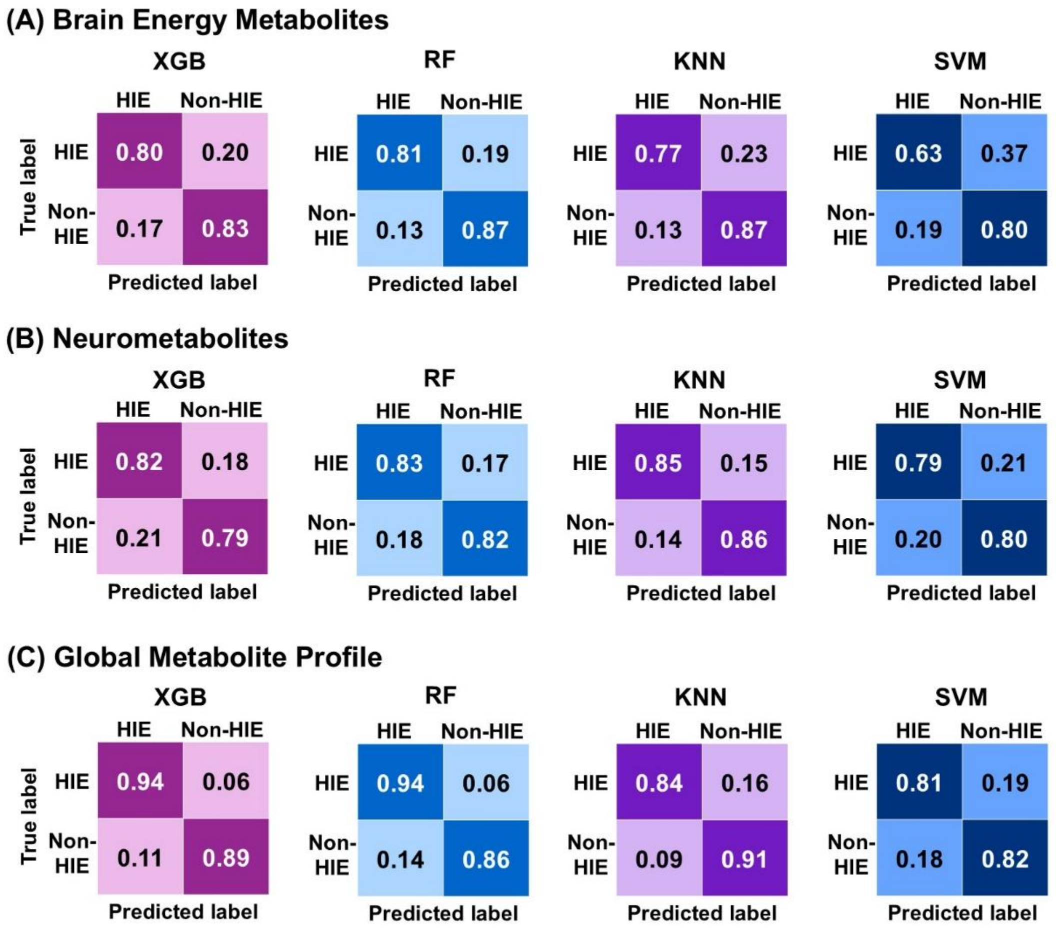


**Figure S3:** Confusion matrices for machine learning models classifying HIE and non-HIE neonates using metabolite panels. (A) brain energy metabolites, (B) neurometabolites, and (C) global metabolite profile. Each matrix depicts the proportion of subjects with HIE and non-HIE classification.

**Table S1:** This table represents plasma metabolite concentrations under two post-collection storage conditions. Blood samples stored at room temperature (~25 °C) for 0-8h show progressive, time-dependent alterations in multiple metabolites, indicating loss of metabolic integrity. In contrast, samples placed on ice (~0-4 °C) immediately after collection and maintained for 0-8h exhibit largely stable metabolite concentrations, consistent with preserved metabolic integrity over the storage interval.

| **Room Temperature (~25°C)** | | | | | |
|---|---|---|---|---|---|
| **Metabolites** | **0hr** | **2hr** | **4hr** | **6hr** | **8hr** |
| Lactate | 0.92 | 1.50 | 2.02 | 2.74 | 3.07 |
| Glucose | 2.24 | 1.71 | 1.42 | 1.09 | 0.83 |
| Glutamine | 0.32 | 0.30 | 0.28 | 0.28 | 0.07 |
| **Ice (~0°C - 4°C)** | | | | | |
| **Metabolites** | **0hr** | **2hr** | **4hr** | **6hr** | **8hr** |
| Lactate | 0.92 | 0.92 | 1.03 | 1.11 | 1.19 |
| Glucose | 2.24 | 2.24 | 2.13 | 2.06 | 2.17 |
| Glutamine | 0.32 | 0.33 | 0.32 | 0.34 | 0.34 |

**Table S2:** This table represents the cross-validation results for Partial Least Squares Discriminant Analysis (PLS-DA) applied to NMR spectroscopy data from HIE and non-HIE neonatal blood samples. Table contains performance measures across different component numbers (1 to 5), including Accuracy, $R^2Y$ (explained variance in class label), $R^2X$ (explained variance in metabolites) and $Q^2$ (predictive variance). These metrics demonstrate effectiveness of the model in differentiating sample groups as components are added, with optimal accuracy and variance achieved with higher component numbers.

| Components | Accuracy | $R^2Y$ | $R^2X$ | $Q^2$ |
|---|---|---|---|---|
| 1 | 0.78 | 0.35 | 0.46 | 0.78 |
| 2 | 0.86 | 0.44 | 0.66 | 0.89 |
| 3 | 0.85 | 0.50 | 0.72 | 0.83 |
| 4 | 0.81 | 0.51 | 0.76 | 0.80 |
| 5 | 0.83 | 0.51 | 0.81 | 0.80 |
| **Average** | **0.83** | **0.46** | **0.68** | **0.82** |

**Table S3:** This table represents significantly altered plasma metabolites distinguishing HIE from non-HIE neonates based on volcano plot analysis. This table summarizes list of log2 fold change ($log_2FC$), p-values, and $-log_{10}$(p-values) quantified for each metabolite through volcano plot. Positive $log_2FC$ values indicate higher metabolite abundance in HIE neonates relative to non-HIE controls, while negative $log_2FC$ values denote downregulation of metabolites in HIE compared to non-HIE.

| Metabolite | log2 (FC) | p-value | $log_{10}$ (p-value) | Significance |
|---|---|---|---|---|
| Lactate | 0.778 | 4.35E-07 | 6.362 | NS |
| Glutamine | -1.279 | 1.50E-08 | 7.824 | **Significant** |
| Succinate | 1.280 | 4.68E-02 | 1.329 | **Significant** |
| NADH | -0.550 | 3.51E-02 | 1.455 | NS |
| Glucose | -1.318 | 2.72E-03 | 2.565 | **Significant** |
| Creatine Phosphate | -0.248 | 3.46E-01 | 0.461 | NS |
| N-acetylaspartate | 0.321 | 3.12E-01 | 0.505 | NS |
| Myo-Inositol | 0.106 | 7.22E-01 | 0.142 | NS |
| 2-Hydroxybutyrate | -0.401 | 6.42E-02 | 1.193 | NS |
| Choline | 0.541 | 3.56E-02 | 1.448 | NS |
| Glycerol | 1.342 | 1.66E-01 | 0.780 | NS |
| Histamine | -0.225 | 1.66E-01 | 0.779 | NS |
| Phenylalanine | 0.007 | 9.65E-01 | 0.016 | NS |
| Glycine | 0.406 | 1.26E-01 | 0.900 | NS |
| Valine | -0.031 | 8.71E-01 | 0.060 | NS |
| Alanine | 0.489 | 2.22E-02 | 1.653 | NS |
| Glutamate | 1.026 | 8.06E-04 | 3.094 | **Significant** |
| Acetate | 0.445 | 8.71E-02 | 1.060 | NS |
| Pyruvate | 0.133 | 8.30E-01 | 0.081 | NS |
| Creatine | 0.347 | 3.79E-01 | 0.421 | NS |
| Tyrosine | 0.296 | 1.84E-01 | 0.734 | NS |
| Taurine | 0.735 | 1.04E-02 | 1.982 | NS |
| Pyroglutamate | 3.884 | 9.77E-05 | 4.010 | **Significant** |
| Histamine | -0.225 | 1.66E-01 | 0.779 | NS |

Abbreviations: FC, Fold Change; NS, Not Significant.

**Table S4:** Significantly altered plasma metabolites discriminating HIE from non-HIE neonates based on one-way ANOVA. The table lists metabolites ranked by F-statistic, together with corresponding t-statistics from pairwise group comparisons and associated p-values. The -$\log_{10}$(p-value) column reflects the strength of statistical significance, with higher values indicating more robust differences between groups. Metabolites such as lactate, glutamine, glutamate, and pyroglutamate exhibit the largest F and t values and the lowest p-values, highlighting major perturbations in glycolysis, TCA cycle activity, and glutamate metabolism.

| Feature | F-stat | t-stat | -$\log_{10}$(p-value) |
|---|---|---|---|
| Glutamine | 39.83 | 6.23 | 1.50E-08 |
| Lactate | 30.36 | -5.62 | 4.35E-07 |
| Pyroglutamate | 16.85 | -4.25 | 9.77E-05 |
| Glutamate | 12.14 | -3.55 | 8.06E-04 |

**Table S5:** Metabolic Pathway Enrichment Analysis in HIE Neonates. This table presents the results of metabolic pathway enrichment analysis performed on significantly altered plasma metabolites identified in neonates with birth/perinatal induced hypoxic-ischemic encephalopathy (HIE). For each pathway, the number of matched metabolites (Hits), the enrichment score, and the statistical significance expressed as $-\log_{10}$(p-value) are listed. Higher enrichment scores and $\log_{10}$(p) values indicate stronger overrepresentation of the pathway among the altered metabolites. The results highlight key metabolic disturbances involving energy metabolism, amino acid turnover, redox balance, and mitochondrial function in early HIE.

| **Pathways** | **Hits** | **Enrichment Score** | **$-\log_{10}$(p)** |
|---|---|---|---|
| Warburg Effect | 8 | 7.843 | 5.726 |
| Urea Cycle | 6 | 11.928 | 5.397 |
| Glutamate Metabolism | 7 | 8.121 | 5.100 |
| Arginine and Proline Metabolism | 7 | 7.495 | 4.857 |
| Glucose-Alanine Cycle | 4 | 17.094 | 4.325 |
| Alanine Metabolism | 4 | 13.115 | 3.821 |
| Gluconeogenesis | 5 | 8.432 | 3.747 |
| Aspartate Metabolism | 5 | 7.949 | 3.620 |
| Glutathione Metabolism | 4 | 11.142 | 3.529 |
| Transfer of Acetyl Groups into Mitochondria | 4 | 10.127 | 3.360 |
| Malate-Aspartate Shuttle | 3 | 16.667 | 3.267 |
| Pyruvate Metabolism | 5 | 5.924 | 3.003 |
| Ammonia Recycling | 4 | 7.181 | 2.770 |
| Citric Acid Cycle | 4 | 6.957 | 2.717 |
| Amino Sugar Metabolism | 4 | 6.745 | 2.666 |

**Table S6:** Characteristics of ML Models for different panels of significant metabolites for classifying HIE and non-HIE.

| Panels | Models | Sensitivity (%) | Specificity (%) | BA (%) | ROC-AUC |
|---|---|---|---|---|---|
| **Energy Metabolites** (Lactate, Glucose, Succinate, Alanine) | XGB | 83.0 ±1 4.7 | 79.3 ± 9.5 | 81.2 ± 8.6 | 0.86 ± 0.07 |
| | RF | 86.5 ± 12.3 | 81.6 ± 9.6 | 83.7 ± 7.6 | 0.90 ± 0.07 |
| | KNN | 80.7 ± 14.5 | 76.5 ± 7.5 | 78.7 ± 8.1 | 0.88 ± 0.07 |
| | SVM | 80.1 ± 18.7 | 67.3 ± 9.7 | 71.5 ± 11.0 | 0.77 ± 0.11 |
| **Neurometabolites/ Neurochemicals** (Glutamine, Glutamate, Choline, Taurine) | XGB | 79.2 ± 16.7 | 81.1 ± 10.0 | 80.8 ± 9.7 | 0.89 ± 0.10 |
| | RF | 81.6 ± 16.5 | 84.0 ± 13.4 | 82.4 ± 10.3 | 0.92 ± 0.09 |
| | KNN | 85.7 ± 12.8 | 86.2 ± 13.0 | 85.2 ± 8.9 | 0.94 ± 0.07 |
| | SVM | 80.0 ± 15.9 | 78.5 ± 13.0 | 79.4 ± 11.7 | 0.85 ± 0.11 |
| **Global Metabolite Profile** (Lactate, Glucose, Succinate, Alanine, Glutamine, Glutamate, Choline, Taurine, Pyroglutamate) | XGB | 88.6 ± 12.7 | 93.8 ± 7.5 | 91.2 ± 6.9 | 0.97 ± 0.03 |
| | RF | 85.9 ± 11.6 | 94.2 ± 7.6 | 90.1 ± 5.9 | 0.97 ± 0.03 |
| | KNN | 91.2 ± 9.9 | 85.6 ± 8.7 | 87.7 ± 5.6 | 0.95 ± 0.04 |
| | SVM | 81.9 ± 12.0 | 80.8 ± 9.7 | 81.5 ± 7.5 | 0.90 ± 0.06 |

Abbreviations: BA, Balanced Accuracy; ROC-AUC, receiver operating characteristic - area under the curve.
